# Microstructural asymmetry in the human cortex

**DOI:** 10.1101/2024.04.08.587194

**Authors:** Bin Wan, Amin Saberi, Casey Paquola, H. Lina Schaare, Meike D. Hettwer, Jessica Royer, Alexandra John, Lena Dorfschmidt, Şeyma Bayrak, Richard A.I. Bethlehem, Simon B. Eickhoff, Boris C. Bernhardt, Sofie L. Valk

## Abstract

While macroscale brain asymmetry and its relevance for human cognitive function have been consistently shown, the underlying neurobiological signatures remain an open question. Here, we probe layer-specific microstructural asymmetry of the human cortex using intensity profiles from *post-mortem* cytoarchitecture. An anterior-posterior cortical pattern of left-right asymmetry was found, varying across cortical layers. A similar anterior-posterior pattern was observed using *in vivo* microstructural imaging, with *in vivo* asymmetry showing the strongest similarity with layer III. Microstructural asymmetry varied as a function of age and sex and was found to be heritable. Moreover, asymmetry in microstructure corresponded to asymmetry of intrinsic function, in particular in sensory areas. Last, probing the behavioral relevance, we found differential association of language and markers of mental health with asymmetry, illustrating a functional divergence between inferior-superior and anterior-posterior microstructural axes anchored in microstructural development. Our study highlights the layer-based patterning of microstructural asymmetry of the human cortex and its functional relevance.

## Main

Hemispheric specialization is a crucial aspect of brain organization that supports human cognitive functions, including language and attention (Hartwigsen et al., 2021; Hervé et al., 2013; Ocklenburg & Mundorf, 2022; Toga & Thompson, 2003; Vallortigara & Rogers, 2020; Wan et al., 2022). Previous research has shown neuroanatomical differences between the left and right hemispheres at the macroscale (Amunts et al., 2003; Glasser et al., 2022; Goldberg et al., 2013; Kong et al., 2018; Kuo & Massoud, 2022; Sha et al., 2021; Williams et al., 2023). Specifically, cortical thickness exhibits an asymmetrical pattern that extends from anterior (left) to posterior (right) regions (Kong et al., 2018; Liao et al., 2023). These structural differences between the left and right hemispheres of the brain may have underlying genetic components, as suggested by (Sha et al., 2021). However, the meso-and micro-architectural basis of the observed asymmetry at the macroscopic level and its potential functional impact are not well understood. In this study, we used a multilevel approach to address this question, detailing structural asymmetry in a histological *post-mortem* brain map and three *in vivo* samples.

The neocortex is composed of six layers that contain neurons of varying size and density. These layers, arranged from the pial to gray-white matter boundary, consist of layer I, which is rich in apical dendrites and axon terminals, layers II and III, rich in pyramidal cells, layer IV, with densely packed neurons, layer V, containing small (5a) or large (5b) pyramidal neurons, and layer VI, featuring corticothalamic pyramidal cells (Amunts et al., 2013; García-Cabezas et al., 2019; Zhang & Deschênes, 1997). It is important to note that laminar and cytoarchitectonic features, which are crucial in qualitative studies, vary across the cortex. For instance, while sensory regions exhibit a well-laminated structure and high cell density, association areas, including language networks, display a reduced cell density and less distinct laminar structure (Amunts et al., 2013; García-Cabezas et al., 2020; Paquola et al., 2020, 2022). Although there is limited evidence of asymmetry in cortical cytoarchitecture, understanding this phenomenon could be crucial for advancing our knowledge of brain function. For instance, a *post-mortem* study that focused on a classic language region, the inferior frontal gyrus, reported a higher cell density in the left hemisphere compared to the right hemisphere (Amunts et al., 2003).

Although *post-mortem* data can provide new insights of cortical microstructure and its potential asymmetry at the microscale, it cannot be linked directly to individual differences and potential functional relevance. Recent advancements in quantitative magnetic resonance imaging (MRI) have made it possible to obtain detailed region-wise *in vivo* microstructure information based on imaging markers such as T1w/T2w (Glasser et al., 2022; Glasser & Essen, 2011), quantitative T1 (qT1) relaxometry (Nieuwenhuys, 2013; Royer et al., 2022; Sereno et al., 2013; Shams et al., 2019), and magnetization transfer (MT) (Duyn, 2018; Harrison et al., 2015; Sled, 2018). *In vivo* quantitative MRI captures the higher intensity in sensory areas and lower intensity in transmodal dys-and agranular cortical regions (Paquola et al., 2022; Paquola, Wael, et al., 2019). Such a differentiation between sensory and transmodal regions is also present in intrinsic functional organization (Margulies, 2016), suggesting a common principle of brain organization for microstructure and function (Paquola, Wael, et al., 2019). Such a principle would be in line with the structural model, posing that regions with similar microstructure may be functionally connected (García-Cabezas et al., 2019). Indeed, various studies have demonstrated asymmetry along this sensory-transmodal-axis for intrinsic function (Gonzalez Alam et al., 2022; Labache et al., 2023; Liang et al., 2021; Wan et al., 2022, 2023).

In the current work, we aimed to probe the microstructural basis of cortical asymmetry using a multiscale approach based on high-resolution histology and imaging data. First, we studied the BigBrain, which is an ultra high-resolution whole-brain *post-mortem* histological atlas of a 65-year-old man. It allowed the quantification of asymmetry in cortical cytoarchitecture at the level of individual layers (Amunts et al., 2013; Saberi et al., 2023; Wagstyl et al., 2018, 2020). Second, we studied *in vivo* microstructural asymmetry to evaluate inter-individual variation. For *in vivo* microstructural maps we used the T1w/T2w ratio from 1101 individual images from the Human Connectome Project (HCP) from young adults (Glasser & Essen, 2011; Van Essen et al., 2012). Then, we investigated the asymmetric microstructure-function coupling. Structural brain asymmetry is linked to behavioral differences such as variability in language skills (Fan et al., 2023; Powell et al., 2006; Qi et al., 2019) and mental health (Kong et al., 2022) such as autism (Postema et al., 2019; Sha et al., 2022), attention-deficit/hyperactivity disorder (Postema et al., 2021), schizophrenia (Schijven et al., 2023), and substance dependence (Cao et al., 2021). Finally, based on the healthy HCP sample, we investigated the associations between microstructural asymmetry and individual differences in language skills, as well as its potential relevance to mental health traits including depression, anxiety, somatic, avoidant, ADHD, and antisocial phenotypes. Given that different imaging sequences have been proposed to measure microstructure *in vivo,* as mentioned above, we leveraged these measures to verify our results including qT1 relaxometry from a dataset (n = 50) for microstructure-informed connectomics (MICs) in young adults (Royer et al., 2022) and MT maps from a longitudinal cohort of adolescents and young adults (n = 286) acquired as part of the Neuroscience In Psychiatry Network (NSPN).

## Results

### Differentiable patterns of microstructural asymmetry as a function of cortical depth in a ultra-high resolution post-mortem sample (Figure 1)

We first mapped the cortical cytoarchitecture asymmetry using ultra-high resolution *post-mortem* data based on the BigBrain (Amunts et al., 2013, 2020). The sliced sections (20 µm) of the BigBrain were cell-body stained, scanned and reconstructed in 3D, resulting in an ultra-high resolution atlas (100 µm^3^) (**Figure 1A**). Using the cortical cell-staining intensity of BigBrain as a feature, a six-layer cortical segmentation (60 surfaces) was obtained of the whole cerebral cortex via a convolutional neural network algorithm (Wagstyl et al., 2020). Multimodal (Glasser et al., 2016) and Cole-Anticevic (CA) parcellation (Ji et al., 2019) were utilized to downsample the maps into 360 regions and 12 networks. The CA networks consisted of primary visual (Vis1), secondary visual (Vis2), somatomotor (SMN), cingulo-opercular (CON), dorsal attention (DAN), language (LAN), frontoparietal (FPN), auditory network (AUD), default mode (DMN), posterior multimodal (PMN), ventral multimodal (VMN), and orbito-affective (OAN). To prevent measurement bias, the mean intensity for layer profiles was regressed out separately for the left and right hemispheres (**Figure 1B**).

Figure 1C and **Supplementary Figure S1** show the mean residual intensity maps. The left-right asymmetry index (AI) was calculated by subtracting the right hemisphere from the left hemisphere. The overall mean map (averaged across layers) showed left-right AI from the anterior to posterior direction (Figure 1D), indicating that the left hemisphere showed higher cell staining intensity in anterior regions but lower intensity in posterior regions, compared to the right hemisphere. The rightward anchor (most rightward region) was located at the AUD and the leftward anchor was located at the LAN at the network level. The LAN also exhibited strong leftward asymmetry in superficial layers, but became rightward asymmetric from layer III onwards, with a peak in layers IV and VI (Figure 1E). Regarding the entire cortex, superficial layers showed anterior-posterior asymmetry and deep layers showed inferior-superior asymmetry (**Supplementary Figure S1**). To summarize the more left-or right-ward asymmetry along six layers (60 surfaces), we calculated the skewness of AI (Figure 1F). Skewness differentiation was observed between more leftward asymmetry in somatomotor and more rightward asymmetry in auditory networks.

**Figure 1.**
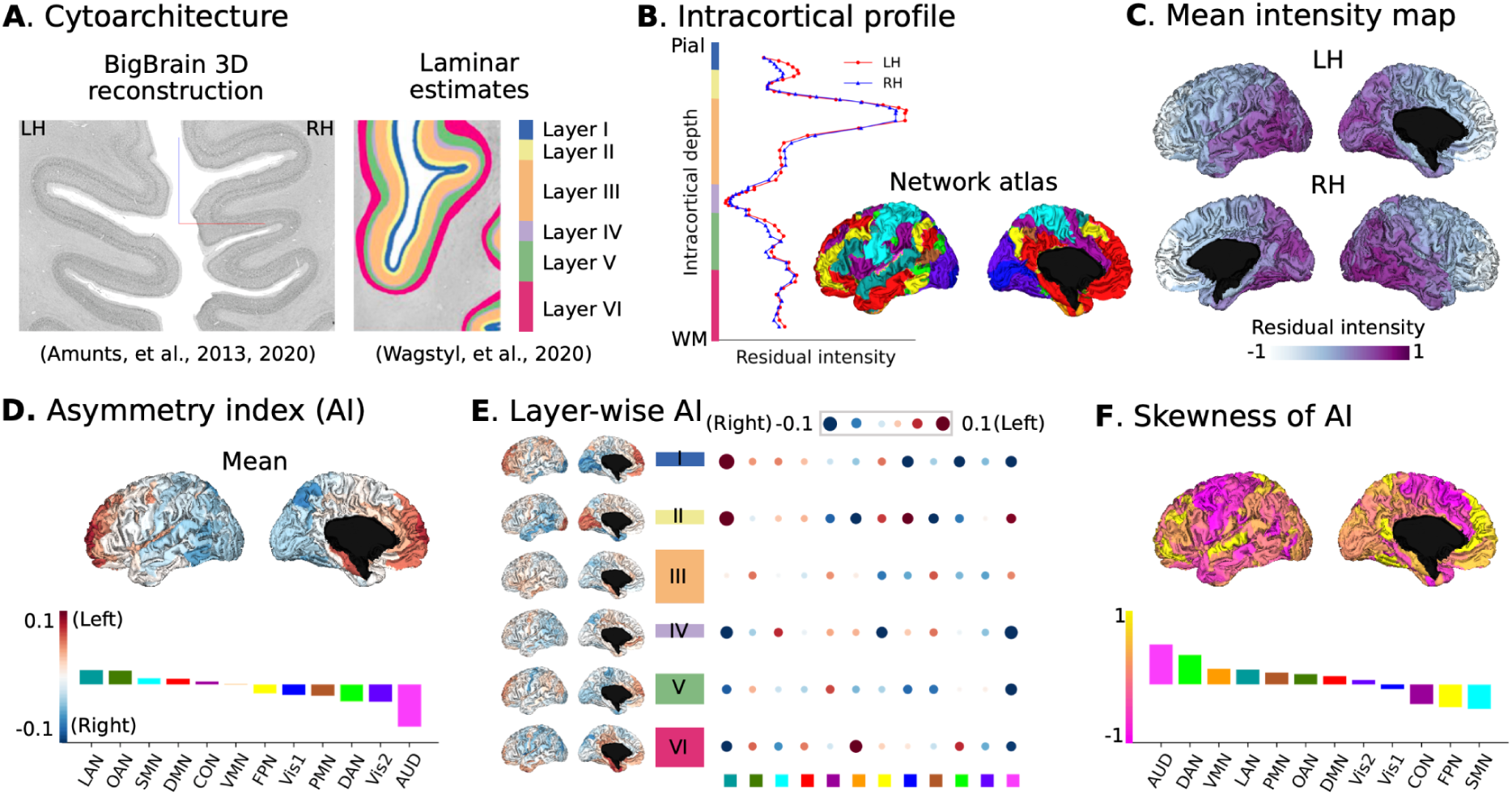
Microstructural asymmetry in BigBrain using cytoarchitecture. **A.** BigBrain 3D histological reconstruction (Amunts et al., 2013, 2020) and six-layer estimates (Wagstyl et al., 2020). **B.** Intracortical staining intensity profiles. Red and blue lines indicate the left and right hemispheres, respectively. **C.** Mean intensity maps across 6 layers. **D.** Mean asymmetry across layers. Red and blue indicate asymmetry index (AI) left > right and right > left. **E.** Layer-wise AI for Bigbrain: six-layer parcel-wise AI brain maps and network-wise heatmap. **F.** Skewness map across asymmetry along 60 points of intracortical depth. Atlas-defined networks include primary visual (Vis1), secondary visual (Vis2), somatomotor (SMN), cingulo-opercular (CON), dorsal attention (DAN), language (LAN), frontoparietal (FPN), auditory network (AUD), default mode (DMN), posterior multimodal (PMN), ventral multimodal (VMN), orbito-affective (OAN).

### Translation to microstructure-sensitive in vivo MRI (Figure 2)

After establishing asymmetry in cytoarchitecture using a post-mortem sample, we aimed to extend the work by using *in vivo* proxies to capture microstructural differentiation in the cortex. Specifically, we extracted intensity data from HCP T1w/T2w maps (n = 1101) and summarized them using a multimodal parcellation and the CA network atlas. T1w/T2w intensity ranges were homogenized by z-scoring intensity values, vertex-wise, independently for each hemisphere (Figure 2A).

In this *in vivo* sample, we observed a populational asymmetry (using Cohen’s d) pattern from anterior to posterior (Figure 2B). At the network level, the rightward effect anchors were located at Vis2 (Cohen’s d = -2.57, *P*_FDR_ < 0.001) and DAN (Cohen’s d = -2.19, *P*_FDR_ < 0.001); the leftward effect anchors were situated at FPN (Cohen’s d = 2.53, *P*_FDR_ < 0.001) and LAN (Cohen’s d = 2.13, *P*_FDR_ < 0.001). For networks’ effect sizes of the left-right asymmetry see **Supplementary Table S1**. Additionally, we observed a significant spatial correlation between the BigBrain and mean HCP asymmetry maps (*r* = 0.482, *P*_variogram_ = 0.007, Figure 2C). This suggests a similarity in spatial patterns between *post-mortem* cytoarchitectural and *in vivo* microstructure asymmetry. Upon further analysis of each layer, we found that in particular layer III exhibited a strong similarity between the two asymmetry maps (*r* = 0.513, *P*_variogram_ < 0.001). Overall, significant correlations were situated in layer I-IV, but not in layer V and VI.

Following this, and building on the twin-pedigree design of the HCP sample, we calculated the heritability (*h*^2^) of asymmetry (Figure 2D). The three most heritable networks were: Vis2 (*h*^2^ = 0.51, SE = 0.05, *P*_FDR_ < 0.001), DAN (*h*^2^ = 0.43, SE = 0.05, *P*_FDR_ < 0.001), and FPN (*h*^2^ = 0.39, SE = 0.06, *P*_FDR_ < 0.001), see **Supplementary Table S2**. The spatial correlation between absolute AI score and heritability maps was Pearson *r* = 0.469 with *P*_variogram_ = 0.001, indicating that regions that are more asymmetric are also more heritable.

Last, to probe potential markers of individual variation, we studied the effects of sex, based on a self-reported question of assigned sex at birth, and age on microstructural asymmetry (Figure 2E). Both the sex and age *t*-maps revealed an anterior to posterior direction and were correlated with the mean *in vivo* AI map (*r*_sex_ = 0.961, *P*_variogram_ < 0.001; *r*_age_ = 0.690, *P*_variogram_ < 0.001). These findings suggest that microstructural intensity is more asymmetric in males and younger individuals, in this relatively young sample. Details for comparisons in functional networks are shown in Figure 2F and G. For convenient visualization, we divided age into two groups (i.e., > 28 years and < 29 years) but t-values were reported based on continuous age. There were 11 out of 12 networks statistically significant after FDR correction for sex comparisons, excluding only DMN (*t* = -0.422, *P*_FDR_ = 0.673). There were 4 out of 12 networks statistically significant for age comparisons including Vis2 (*t* = -2.797, *P*_FDR_ = 0.031), DAN (*t* = -2.589, *P*_FDR_ = 0.039), FPN (*t* = 4.826, *P*_FDR_ < 0.001), and LAN (*t* = 2.456, *P*_FDR_ = 0.042).

**Figure 2.**
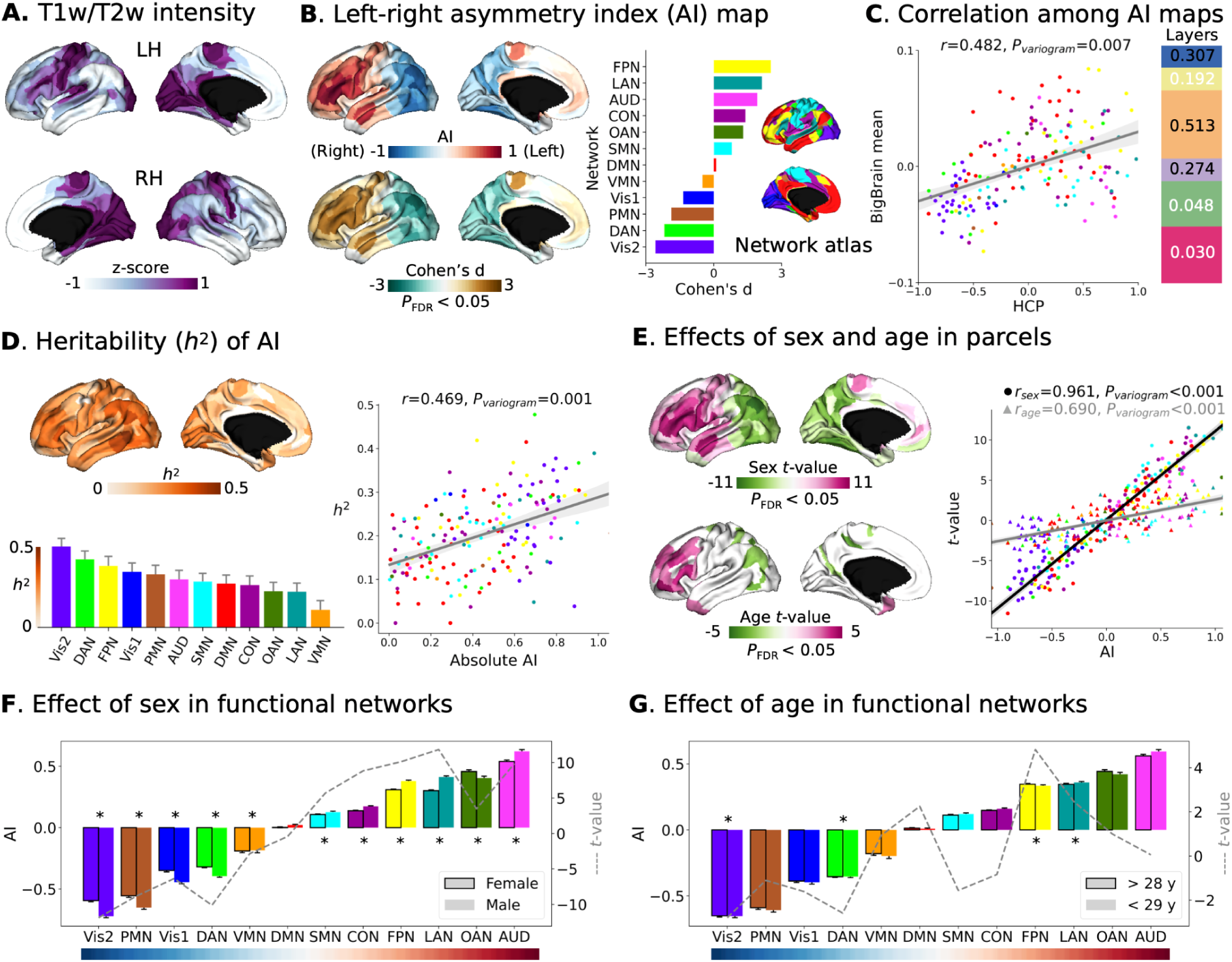
Microstructural asymmetry in Human Connectome Project (HCP) using T1w/T2w images. **A**. T1w/T2w intensity values for left and right hemispheres (Z-scored separately). Deeper purple indicates higher intensity. **B.** The mean asymmetry index (AI) and related Cohen’s d maps calculated across subjects. Red/brown and blue/green indicate left-and right-ward asymmetry direction at populational level. AI was also summarized into functional networks with mean and standard error in the barplot. Cohen’s d map was thresholded at *P*_FDR_ < 0.05. **C**. Spatial correlation between mean HCP AI and BigBrain AI maps. A variogram permutation test was used to account for the spatial autocorrelation. Black and white values indicate significance after variogram permutation at *P* < 0.05 level. **D**. Heritability map and network barplot estimated by individual variation of AI. The heritability map was thresholded at *P*_FDR_ < 0.05. Right panel is the spatial correlation between mean AI and heritability maps. *T*-maps of sex and age effects in the model of AI = 1 + sex + age. Purple red indicates higher leftward asymmetry in females and in older people, respectively. Right panel is the spatial correlation between mean AI and t-value maps. Round and triangle dots represent sex and age. **E**. *T*-maps of sex and age effects in the model of AI = 1 + sex + age. Purple red indicates higher leftward asymmetry in females and in older people, respectively. Right panel is the spatial correlation between mean AI and t-value maps. Round and triangle dots represent sex and age. **F** and **G** plot the detailed sex and age effects in functional networks (AI mean and standard errors). Dashed lines indicate *t*-value for sex and age, respectively. * indicates statistical significance after FDR correction. The colors of dots and bars in all plots reflect atlas-defined functional networks including primary visual (Vis1), secondary visual (Vis2), somatomotor (SMN), cingulo-opercular (CON), dorsal attention (DAN), language (LAN), frontoparietal (FPN), auditory network (AUD), default mode (DMN), posterior multimodal (PMN), ventral multimodal (VMN), orbito-affective (OAN).

### Microstructural asymmetry is linked with asymmetry in intrinsic function (Figure 3)

After establishing microstructural asymmetry in both *post-mortem* and *in-vivo* markers, we investigated its functional association. To achieve this, we utilized resting state functional connectivity (FC) in the same sample (i.e., HCP), as in previous research (Wan et al., 2022).

To investigate the relationship between microstructure and function asymmetry at the group level, we divided the mean microstructural asymmetry map into 10 bins (see Figure 3A-i). We then calculated the average functional connectivity (FC) within each bin (see Figure 3A-ii) and determined the functional connectivity asymmetry by subtracting the right hemisphere (RH) from the left hemisphere (LH) and dividing by the sum of RH and LH. Our analysis focused on intra-hemispheric (i.e., LH_LH and RH_RH) connectivity. Functional connectivity was observed to be stronger between regions that exhibited similar patterns of asymmetry, compared to those with varying degrees of asymmetry (Figure 3A-iii). This relationship was quantitatively assessed by calculating region-wise microstructure-function correlation between T1w/T2w AI map and FC AI profile (Figure 3B-i). We found coupling was strongest in central and superior temporal areas and weakest in prefrontal and parietal areas, both at the group level and individual level (*r* = 0.664, *P*_variogram_ < 0.001). As demonstrated in the map of mean and standard deviation, this coupling exhibited significant individual variability, particularly in areas of high coupling (Figure 3B-ii).

Finally, to study how asymmetry in microstructure-function coupling varied between regions, we calculated the individual co-variation between microstructural and FC asymmetry per parcel across subjects (Figure 3C-i). Then we extracted the top 10% of the co-variation to calculate the affinity matrix in a normalized angle. Finally, using principal component analysis (PCA), we decomposed the affinity matrix by rows and columns. Row PCs summarized microstructural similarity in functional profiles and column PCs summarized functional similarity in microstructure (Figure 3C-ii). For the microstructural PCs, the first two components accounted for 26.6% and 17.4% of the total variance, respectively. PC1 displayed a dissimilarity axis from the dorsolateral prefrontal to the precentral gyrus, while PC2 displayed a dissimilarity axis from the temporoparietal junction to the lateral prefrontal areas. Regarding the functional PCs, the first two components accounted for 48.4% and 25.8% of the total variance, respectively. PC1 differentiated prefrontal from visual regions, and PC2 differentiated sensory from association regions (Figure 3C-iii).

**Figure 3.**
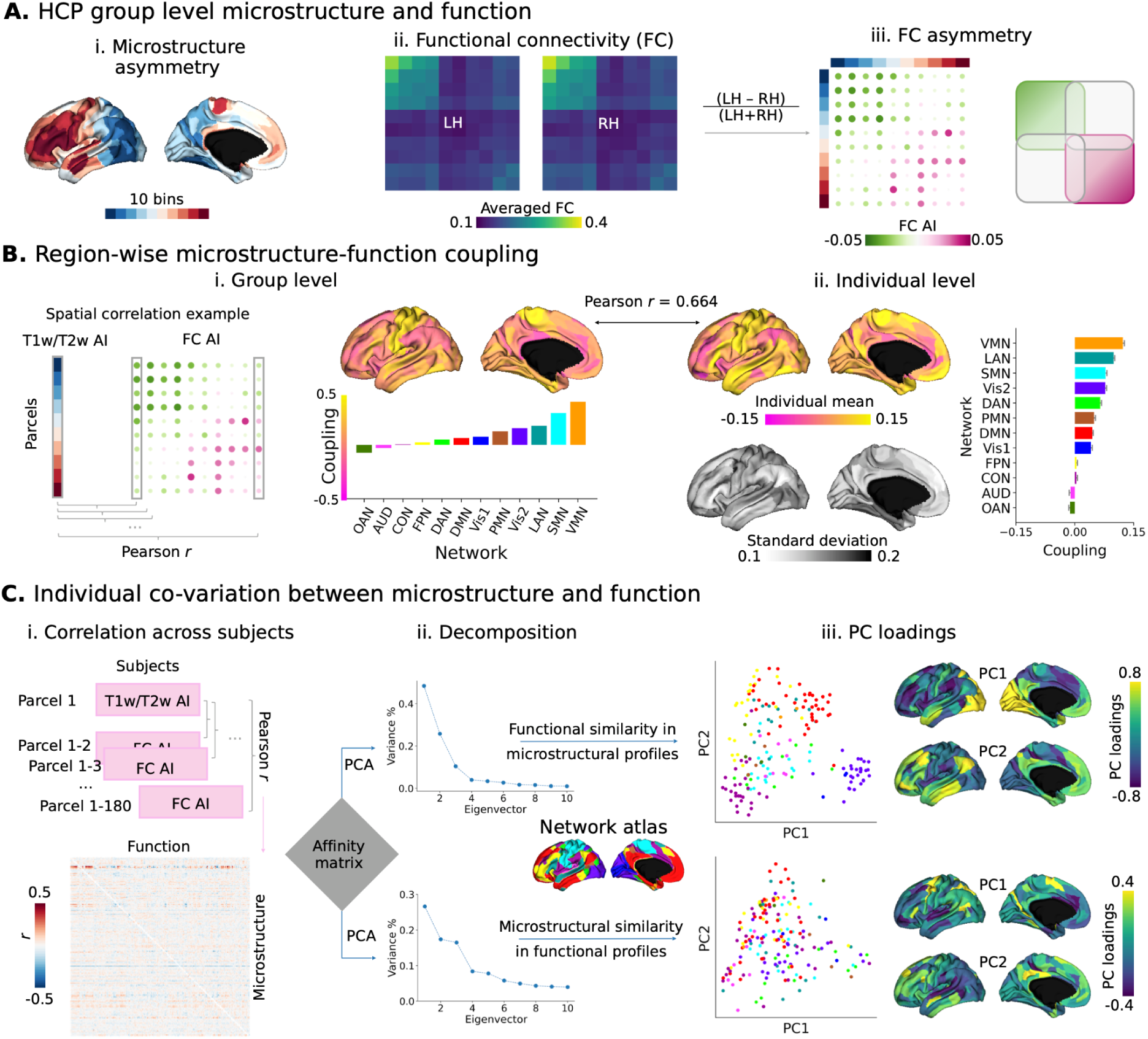
Microstructure-function relationship in asymmetry using HCP T1w/T2w and resting state functional images. **A**-i. 10 bins (18 parcels per bin) categorized from **Figure 2B** mean AI map (T1w/T2w). **A**-ii. Group-level resting state functional connectivity (FC) matrix averaged by bins. **A**-iii. FC asymmetry calculated by (LH - RH)/(LH + RH) sorted by bins. Purple-red and green indicate left-and right-ward asymmetry. Scatters are colored by functional networks. **B**. Region-wise microstructure-function coupling was calculated by Pearson correlation coefficient between 180 parcels of T1w/T2w and FC AI per column. Left panel shows coupling between mean maps at the group level (i) and the right panel shows mean and standard deviation of coupling (ii). **C.** Individual covariation between microstructure and function. Matrix in (i) represents the Pearson r between parcel T1w/T2w AI and FC AI across subjects. Then, the parcel-wise affinity matrix was computed and principal component analysis (PCA) was employed to decompose the matrix to detect the inter-region similarity axes (ii). Upper and lower panels are microstructural and functional decomposition (iii). The first two eigenvectors and eigenvalues (PC loadings) are plotted with similar colors in ‘viridis’ indicating similar profiles between regions. Atlas-defined networks include primary visual (Vis1), secondary visual (Vis2), somatomotor (SMN), cingulo-opercular (CON), dorsal attention (DAN), language (LAN), frontoparietal (FPN), auditory network (AUD), default mode (DMN), posterior multimodal (PMN), ventral multimodal (VMN), orbito-affective (OAN).

### Microstructural asymmetry relates to individual variability in language skills and mental health (Figure 4)

Our last aim was to investigate the behavioral relevance of microstructural asymmetry. Language scores were obtained through reading and picture vocabulary tests. Mental health scores were assessed using the Adult Self-Report and DSM-Oriented Scale, which included depression, anxiety, somatic, avoidant, ADHD, and antisocial problems. Therefore, two variables for language, six variables for mental health, and 180 brain variables were included. To achieve this, we conducted a canonical correlation analysis (CCA), which is a multivariate approach to estimate latent dimensions for multiple independent and dependent variables via correlation.

**Figure 4.**
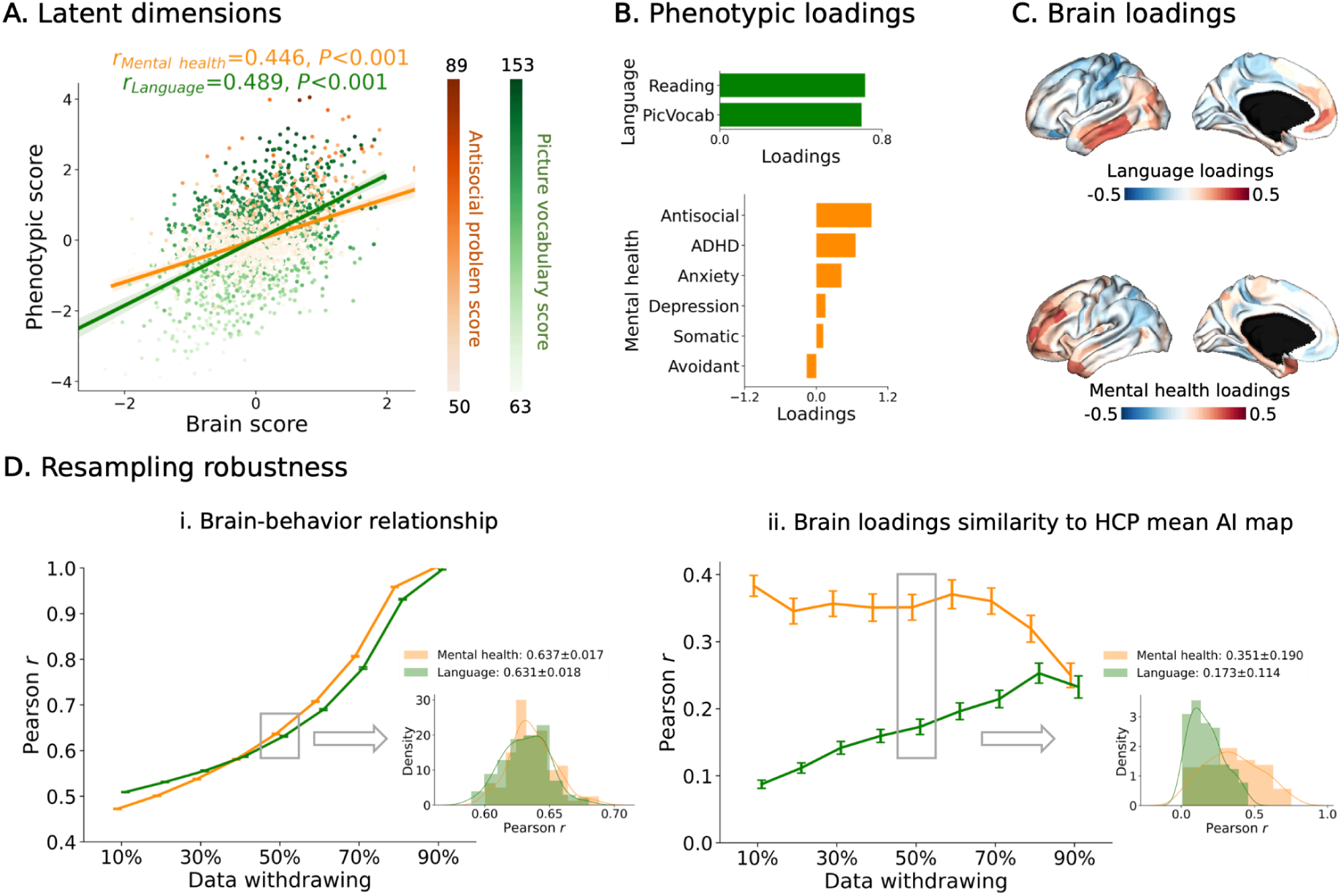
Canonical correlation analysis (CCA) between microstructural asymmetry features and language/mental health in HCP. **A**. Correlation between latent dimensions of the brain and phenotype. Orange and green indicate mental health and language, where the latent dimensions explain antisocial behavior and picture vocabulary scores most. **B**. Phenotypic loadings of the first latent dimension for language and mental health. **C**. Brain loadings of the first latent dimension for language and mental health. **D**. Resampling data to test the performance of CCA. We withdrew data from 10% to 90% by pseudo-randomization using twin classes and resampled 100 times. Mean and standard error bars were shown in the charts. Data withdrawal of 50% was selected to show the distribution across the 100 samples and other percentages see **Supplementary Figure S4.**

We calculated the first latent dimension for language and mental health separately. Statistically significant correlations were found between the first latent dimensions of microstructural asymmetry and behavioral markers (*r*_language_ = 0.489, *P* < 0.001; *r*_mental health_ = 0.446, *P* < 0.001, Figure 4A). Picture vocabulary and antisocial problem scores had the strongest loadings on the respective brain-behavior latent components (Figure 4B). Moreover, we observed a behavioral marker divergence of spatial patterns in brain loadings between language and mental health. Whereas the former showed a differentiation between superior and inferior areas, mental health was linked to differentiation between anterior and posterior portions in asymmetry, with stronger leftward asymmetry in frontal regions associated with reduced mental health (higher scores) (Figure 4C). The anterior-posterior layer-AI maps were similar to mental health brain loadings (*r*_HCP_ = 0.415, *P*_variogram_ = 0.024; *r*_layer_III_ = 0.315, *P*_variogram_ = 0.020, **Supplementary Figure S2**) but the more inferior-superior layer-AI map was similar to language brain loadings (*r*_layer_V_ = 0.210, *P*_variogram_ = 0.061, **Supplementary Figure S3**). This may suggest a reduced leftward asymmetry in the frontal lobe and an increased rightward asymmetry in the occipital lobe for individuals with higher mental health scores.

To test the robustness of the findings, we conducted the pseudo-randomized resampling, 100 times, withdrawing 10%-90% of the data based on the twin label (Figure 4D). We show the average result (50% of data withdrawn) in the main figure and remaining in **Supplementary Figure S4**. We found that the first latent brain and behavioral dimensions remained robust (*r*_language_ = 0.631 ± 0.018, *r*_mental health_ = 0.637 ± 0.017). The correlation between mean AI map and brain loadings were *r*_language_ = 0.173 ± 0.114 and *r*_mental health_ = 0.351 ± 0.190.

### Replication analyses

To replicate our findings using independent samples, we conducted *in vivo* analyses on two external datasets with varying imaging measurements for cortical microstructure. These datasets include MICs in young adults (qT1 relaxometry, n = 50) and the NSPN in adolescents and young adults (MT, baseline n = 286).

Regarding MICs qT1 relaxometry data, the sample demographics resembled HCP data (female: 46%, age centered on 25-35 years old). An anterior-posterior asymmetry pattern was again observed (**Supplementary Figure S5**) with a significant correlation between MICs and the HCP AI map (*r* = 0.548, *P*_variogram_ = 0.004). The spatial pattern of the sex effect was also replicated (*r*_sex_ = 0.731, *P*_variogram_ < 0.001) whereas the age effect was not (*r*_age_ = 0.224, *P*_variogram_ = 0.062). We used MT images using multi-parametric mapping (Weiskopf et al., 2013) from NSPN to address potential transmit field issues associated with T1w/T2w. The sample consisted of individuals aged 14 to 25 years (mean ± SD: 19.1 ± 2.9) with a balanced sex ratio (female: 51%). Again we observed the anterior-posterior asymmetry pattern and found a significant correlation (*r* = 0.369, *P*_variogram_ = 0.047) between HCP and NSPN (**Supplementary Figure S6**). In addition, the spatial pattern of age and sex effects could also be replicated in this younger sample (*r* _sex_ = 0.358, *P*_variogram_ < 0.001; *r* _age_ = 0.323, *P*_variogram_ < 0.001). Together, these analyses suggest that microstructural asymmetry is consistent across different microstructural measures and histology.

## Discussion

Asymmetry in structural and functional brain organization is implicated in key human cognitive functions, including language, and is associated with neuropsychiatric conditions. In this study, we used a multiscale approach to investigate microstructural asymmetry at ultra-high resolution and applied our model to *in vivo* data to examine individual variability and functional relevance. A consistent pattern of left-right asymmetry of microstructure, along the anterior to posterior regions, was observed in an ultra high-resolution *post-mortem* human brain and three *in vivo* samples. Using an ultra-high resolution post mortem model, we found that the asymmetry pattern differed across layers, with superficial layers showing an anterior-posterior pattern and deep layers showing an inferior-superior pattern. Furthermore, utilizing an *in vivo* model of microstructure, we demonstrated that microstructural asymmetry varies with age and self-reported sex and is heritable. Finally, we established the functional relevance of these findings by linking microstructural asymmetry to asymmetry in intrinsic functional connectivity profiles as well as behavioral markers that detail individual variations in language skills and mental health traits. This can be viewed as a proxy of neuropsychiatric risk within the healthy population. Language skills vary along an inferior-superior axis, while mental health traits vary along an anterior-posterior axis, suggesting that mental health may be associated with superficial laminar functions but language may coordinate with deep laminar functions, a divergence possibly anchored in neurodevelopmental trajectories of both patterns. Together, our findings illustrate consistent cortical brain asymmetry in cytoarchitecture and microstructure and its functional consequences.

In the current work we estimated asymmetry of human cortical cytoarchitecture and uncovered an overall pattern of left-to right-ward asymmetry in an anterior to posterior axis. Previous studies on regional cytoarchitecture have indicated that the left hemisphere has a higher neuronal density in inferior frontal cortex, i.e., Brodmann area (BA) 45 (Amunts et al., 2003) and dorsolateral prefrontal cortex, i.e., BA 9. These have been attributed to pyramidal neurons in layer III (Cullen et al., 2006). Leftward asymmetry was also reported in anterior regional torque for Broca’s area and rightward asymmetry was reported in posterior regional torque for occipital visuospatial area (Toga & Thompson, 2003). The anterior-posterior differentiation in asymmetry may be related to neurodevelopmental patterning and cortical maturation. Indeed, whereas posterior regions show early postnatal development, anterior regions mature in adolescence, illustrated by variations of cyto-and myelo-architecture and connectivity (Fornito et al., 2019). Though the current study did not evaluate developmental patterning over time, and the subject (the BigBrain data) was also older than 60 years, it is possible that the asymmetry observed is still a consequence of maturational timing in combination with experience dependent plasticity due to the differential functional role of the left and right hemisphere.

Moreover, we observed notable depth-wise variation with the anterior-posterior asymmetry in upper and inferior-superior asymmetry in deep layers. Overall, maturation patterns of microstructure during childhood have been shown to follow a posterior-anterior pattern (Baum et al., 2022). Previous work has reported divergent maturational profiles in intra-cortical microstructure between sensory and paralimbic areas in adolescence (Paquola, Bethlehem, et al., 2019)). In particular, mid-to-deep layers seem to have a preferential development in adolescents, specifically in uni-and hetero-modal areas spatially corresponding to attention and language regions (Paquola, Bethlehem, et al., 2019). In our work, the language network shifted from leftward (superficial layers) to rightward asymmetry (deep layers), which may be related to the development and maturation of cortico-cortical connections (Broser et al., 2012; Kanno et al., 2018; Matsumoto et al., 2004). The laminar architecture of the cortex has an important role in coordinating functional processes and connectivity between cortical regions and subcortex and cortex (Shipp, 2007). For cortico-cortical or cortico-subcortical pathways, cellular and synaptic architectures differ across layers such that they result in distinct computations at the target projection neurons (D’Souza & Burkhalter, 2017). Though observing asymmetry across layers results in novel hypotheses and perspectives on the neuroanatomical origin of functional specializations in the cortex, indicating asymmetry not only varies spatially along the cortical mantle but also as a function of its depth.

Microstructural asymmetry in the anterior-posterior direction could also be identified using *in vivo* imaging, using T1w/T2w maps, qT1 relaxometry, and MT. Previous work has suggested that the anterior-posterior asymmetry pattern in T1w/T2w is partly generated by the transmit field rather than the microstructure itself (Glasser et al., 2022). Yet, cell staining and MT contrast used in the current study are likely less affected by the transmit field, suggesting that at least a part of the effects may go above and beyond signal noise. Taking advantage of the inter-individual model of *in vivo* data, we probed its relevance to age and sex differences in the samples tested. We found that more asymmetric regions overall show a stronger age effect. Increased cortical symmetry with age has been previously reported in the mid-and old-aged sample of the UK Biobank (Korbmacher et al., 2024) and young adults (Kovalev et al., 2003). Looking ahead, the implementation of a normative model could elucidate the developmental trajectory of microstructural asymmetry across the lifespan, facilitating the identification of individual trajectories (Bethlehem et al., 2022). Previous work has also reported marked sex-differences in various indices of cortical structure (DeCasien et al., 2022; Küchenhoff et al., 2023; Liu et al., 2020; Serio et al., 2023), possibly related to differential sex hormonal expression and physiological markers (Wisniewski, 1998). Indeed, in the current work we observed an overall stronger asymmetry in males relative to females. In related work, reported overall decreases in mean microstructure and increases in skewness were observed in women relative to men, and varied as a function of (self-reported) hormonal status (Küchenhoff et al., 2023). However, in the current work we could only touch upon these neuroendocrine, physiology, and age-related factors shaping microstructural brain asymmetry. Further work including hormonal data and broader age ranges may further reveal potential causes and consequences of the observed associations between age and self-reported sex and brain asymmetry.

We found that microstructure and intrinsic function have similar lateralized directions within hemispheres. In the cerebral cortex, regions with similar cytoarchitecture tend to have similar functional connectivity profiles (García-Cabezas et al., 2019; Paquola, Wael, et al., 2019). We showed that this principle may also hold for asymmetry of microstructure and functional connections. In particular, somatomotor and language-related areas show an increased convergence between functional and microstructural asymmetry profiles. These regions have highly specialized functions, possibly suggesting that functional specialization may be accounted for by cortical asymmetry (Duboc et al., 2015; Hartwigsen et al., 2021; Toga & Thompson, 2003; Witelson & Pallie, 1973). Relatedly, via inter-individual co-variation, we observed that regions within functional networks share more similarity in microstructural asymmetry profiles, again underscoring the link between microstructural asymmetry and intrinsic function. This pattern could be interpreted as suggesting that activity-dependent plasticity in part shapes the microstructural asymmetry observed, ultimately supporting similar microstructural profile asymmetry in functionally connected regions. At the same time, the inverse of the model, i.e. microstructure-guided function, revealed clearer organizational patterns.

Probing functional relevance of microstructural asymmetry in terms of behavioral outcomes, we observed a pattern of inferior-superior differentiation for language. This was mainly linked to a differentiation of temporal lobe and sensorimotor regions, which spatially mirrored the asymmetry pattern in layer V of the BigBrain. Layer V is the main output layer of the cortex and largely relays signals to subcortical structures (Shipp, 2007), yet also shows marked divergence of connection profiles as a function of neuron type (Kim et al., 2015). On top of this, we found that cell density is lower in the left relative to the right hemisphere for the language network in the BigBrain. Thus, the observed patterns possibly relate to a differentiation of cells in deeper cortical layers (temporal and sensorimotor) linked to differential output profiles ultimately leading to behavioral differences in language. Second, we found a behavioral marker of mental health to be associated with anterior-posterior differentiation in asymmetry, a pattern present in superficial layers of the BigBrain, and in overall maturational patterns of microstructure in the cortex (Baum et al., 2022). Various studies have reported associations between microstructure and neuropsychiatric conditions including depression (Baranger et al., 2021; Chen et al., 2022; Sacchet & Gotlib, 2017), compulsivity and impulsivity (Ziegler et al., 2019), and schizophrenia (Flynn et al., 2003; Frangou et al., 2020; Palaniyappan et al., 2019). Through large sample size investigation, it would be possible to study the inter-relationships among asymmetries of cortical brain maturation and neurodevelopmental conditions, extending current work on maturational differentiation of symmetric microstructural patterning and links to disease progression (Flynn et al., 2003; Knowles et al., 2022).

While our research has yielded significant insights into cortical microstructural asymmetry, it is important to address several limitations that warrant clarification. Firstly, though the BigBrain offers unique insights into cortical microstructure at ultra high resolution, and links to our *in vivo* model, further work on ultra-high resolution neuroimaging will aid in also understanding layer-level markers of individual variation. Furthermore, while the observed map of age effects in our young adult sample correlated with the mean asymmetry map, no parcels exhibited significant age effects in the MICs and NSPN datasets. The modest changes observed during adolescence may stem from either small effect sizes or inadequate sample sizes to detect statistical significance. Additionally, it is crucial to acknowledge that the mental health data utilized in this study pertains solely to healthy individuals, and caution should be exercised in extrapolating these findings to clinical samples, where extreme conditions may prevail that were not accounted for in our study.

In conclusion, our investigation employed a multiscale approach to study microstructural asymmetry in the human cortex. We delineated laminar-specific asymmetry in a post-mortem sample and individual asymmetry *in vivo*. Our study contributes to advancing our understanding of cortical asymmetry at the microscale, encompassing depth-wise and inter-regional spatial differentiation, age and sex disparities, behavioral genetics based on twin-modeling, integration with functional connectomics, and associations with behavioral markers of language and mental health. These findings hold implications for elucidating the biological mechanisms underlying cortical asymmetry and its functional relevance in health and disease.

## Methods

Datasets we used in present study are open sources and have been approved by their local research ethics committees. The current research complies with all relevant ethical regulations as set by The Independent Research Ethics Committee at the Medical Faculty of the Heinrich-Heine-University of Duesseldorf (study number 2018–317).

### Datasets & image acquisition and preprocessing

#### BigBrain

BigBrain is a 20 µm^3^ ultra-high-resolution atlas of a *post-mortem* human brain from a 65-year-old male created by digital volumetric reconstruction of Merker-stained sections (https://ftp.bigbrainproject.org/) (Amunts et al., 2013). The six layers of BigBrain’s cerebral cortex were previously segmented using a convolutional neural network and the surface reconstruction of the layer boundaries were available (Wagstyl et al., 2020). We extracted layer-wise cortical profiles of the BigBrain cerebral cortex by sampling the staining intensity of 100µm resolution BigBrain images at 10 equivolumetric surfaces along the depth of each layer (60 surfaces in total). The resulting layer-wise cortical profiles reflect the variation of neuronal size and density along the depth of the six cortical layers at each location.

To reduce the computational demands, we downsampled the images from BigBrain native surface space to Glasser multimodal parcellation (Glasser et al., 2016), a homologous atlas with 180 parcels per hemisphere via BigBrainWarp (Paquola et al., 2021). To enhance the functional annotation we employed the cortical functional network atlas (Ji et al., 2019), which includes 12 networks: primary visual (Vis1), secondary visual (Vis2), somatomotor (SMN), cingulo-opercular (CON), dorsal attention (DAN), language (LAN), frontoparietal (FPN), auditory network (AUD), default mode (DMN), posterior multimodal (PMN), ventral multimodal (VMN), and orbito-affective (OAN).

#### HCP

We used T1w/T2w images from the Human Connectome Project (HCP) S1200 release, which can be downloaded from HCP DB (http://www.humanconnectome.org/). HCP S1200 includes 1206 individuals (656 females) that are made up by genetic-identified and reported 334 MZ twins, 152 DZ twins, and 720 singletons. We included individuals for whom the scans and data had been released after passing the HCP quality control and assurance standards (Glasser et al., 2013; Van Essen et al., 2013). Finally, for genetic analyses we included 1101 healthy subjects with a good quality T1w/T2w image (age: 28.8 ± 3.7 years), of which 54.4% were females and 332 were MZ twins.

MRI data were acquired on the HCP’s custom 3T Siemens Skyra equipped with a 32-channel head coil. Two T1w images with identical parameters were acquired using a 3D-MP-RAGE sequence (0.7 mm isovoxels, matrix = 320 × 320, 256 sagittal slices; TR = 2400 ms, TE = 2.14 ms, TI = 1000 ms, flip angle = 8; iPAT = 2). Two T2w images were acquired using a 3D T2-SPACE sequence with identical geometry (TR = 3200 ms, TE = 565 ms, variable flip angle; iPAT = 2). T1w and T2w scans were acquired on the same day. The pipeline used to obtain the Freesurfer-segmentation is described in detail in a previous article (Glasser et al., 2013). The preprocessing steps included co-registration of T1-and T2-weighted scans, then correcting the T1w and T2w images for B1-bias and some B1+ bias (Glasser et al., 2013; Glasser & Essen, 2011). Preprocessed images were nonlinearly registered to MNI152 space, and segmentation and surface reconstruction performed using FreeSurfer 5.3. T1w images were divided by aligned T2w images to produce a single volumetric T1w/T2w image per subject (Glasser & Essen, 2011). Cortical surfaces were aligned using MSMAll (Robinson et al., 2014, 2018) to the hemisphere-matched conte69 template (Van Essen et al., 2012). Notably, this contrast nullifies inhomogeneities related to receiver coils and increases sensitivity to intracortical myelin.

The intensity values were estimated between pial and white matter surfaces. Previous papers have used this data to generate the equivolumetric profile intensity (Paquola, Wael, et al., 2019; Valk et al., 2022). We downsampled the images from conte69 space to the multimodal atlas. In this study, we averaged the intensity values across the equivolumetric surfaces and z-scored the values for left and right hemispheres separately for each subject.

#### MICs

For replication analysis we used the quantitative T1 images of the openly available MRI dataset for Microstructure-Informed Connectomics (MICs) (Royer et al., 2022), which can be downloaded from the Canadian Open Neuroscience Platform’s data portal (https://portal.conp.ca). The dataset comprises multimodal data of 50 healthy young adults (23 women; 29.54 ± 5.62 years; 47 right-handed) and was collected at the Brain Imaging Centre of the Montreal Neurological Institute and Hospital using a 3T Siemens Magnetom Prisma-Fit and a 64-channel head coil. For the acquisition of the qT1 relaxometry data a 3D magnetization prepared 2 rapid acquisition gradient echoes sequence was used (3D-MP2RAGE; 0.8 mm isotropic voxels, 240 sagittal slices, TR = 5000 ms, TE = 2.9 ms, TI 1 = 940 ms, T1 2 = 2830 ms, flip angle 1 = 4°, flip angle 2 = 5°, iPAT = 3, bandwidth = 270 Hz/px, echo spacing = 7.2 ms, partial Fourier = 6/8) . Two inversion images were combined for qT1 mapping. Based on the varying T1 relaxation time in fatty tissue compared to aqueous tissue (Weiskopf et al., 2021), we here use qT1 as an index for gray matter myelin and hence as a proxy for microstructure. The MRI processing tool *micapipe* (Cruces et al., 2022) was used for data preprocessing and intensity extraction. In short, preprocessing included the background denoising of MP2RAGE, reorientation of the T1W and MP2RAGE, N4 bias correction, and intensity rescaling of the T1W images and the non-linear registration to MNI152 space. Further, the cortical surface reconstruction from native T1w acquisitions was carried out using Freesurfer 7.0. The detailed acquisition protocol and preprocessing are described in their prior data publication (Royer et al., 2022).

#### NSPN

The Neuroscience in Psychiatry (NSPN) cohort generally comprises 2245 adolescents aged 14 to 26 years (mean ± SD age: 19.1 ± 3.0 years, female: 54%). Participants were recruited in Cambridgeshire and north London according to a sampling design that balanced sex, ethnicity, and participant numbers in five age strata (14-15, 16-17, 18-19, 20-21, 22-25). Here, we included 286 individuals (mean ± SD age: 19.1 ± 2.9 years, female: 51%) for whom microstructural neuroimaging data were available.

Magnetization Transfer (MT) data were acquired to approximate myelin content using a multi-parametric mapping (MPM) sequence (Weiskopf et al., 2013) on three identical 3T Siemens MRI Scanners (Magnetom TIM Trio) in Cambridge (2 sites) and London (1 site). A standard 32-channel radio-frequency (RF) receive head coil and RF body coil for transmission were used. MPM included three multi-echo 3D FLASH scans: predominant T1-weighting (repetition time (TR) = 18.7 ms, flip angle = 20°), and predominant proton density (PD) and MT-weighting (TR= 23.7 ms; flip angle = 6°). To achieve MT-weighting, an off-resonance Gaussian-shaped RF pulse (4 ms duration, frequency offset from water resonance = 2 kHz; nominal flip angle = 220°) was applied prior to the excitation. Several gradient echoes were recorded with alternate readout polarity at six equidistant echo durations (TE) between 2.2 and 14.7 ms for MT weighted acquisition. The longitudinal relaxation rate and MT signal are separated by the MT saturation parameter, creating a semi-quantitative measurement that is resistant to relaxation times and field inhomogeneities (Hagiwara et al., 2018; Weiskopf et al., 2013). Other acquisition parameters include: 1 mm isotropic resolution, 176 sagittal partitions, field of view (FOV) = 256×240 mm, matrix = 256×240×176, non-selective RF excitation, RF spoiling phase increment = 50 °, parallel imaging using GRAPPA factor two in phase-encoding (PE) direction (AP), readout bandwidth = 425 Hz/pixel, 6/8 partial Fourier in partition direction. The acquisition time was approximately 25 min. Participants wore ear protection and were instructed to lie still.

Surface reconstruction was carried out based on T1-weighted (T1w) images using Freesurfer 5.3.0. The resulting reconstructions underwent visual inspection. Control points were added to improve segmentations, but scans were excluded in cases of persistently poor quality. The detailed description about this dataset and preprocessing are shown in their previous work (Hettwer et al., 2024; Kiddle et al., 2018; Paquola, Bethlehem, et al., 2019; Weiskopf et al., 2013).

### Asymmetry Index

We calculated the asymmetry index (AI) by subtracting right from left hemispheric values in the homologous regions. As noted, we preprocessed the left and right hemispheres separately by regressing out mean surface intensity for the BigBrain data, then standardized the residual intensity. For HCP, MICs, and NSPN, we obtained the mean cortical intensity map, then z-scored the map for left and right hemispheres separately. For functional connectivity asymmetry, we calculated AI by (LH - RH)/(LH + RH).

### Effects of sex and age

We first used fixed effects estimates for the model: AI = 1 + sex + age + sex*age. We found no significant interaction between age and sex. We then used the non-interaction model: AI = 1 + sex + age to obtain the *t* and *P* values of sex and age. False discovery rate (FDR) was then applied for multiple comparison correction for the sex and age *t* value maps. All the steps were performed in Python with the package BrainStat (Larivière et al., 2023). Regions colored in the HCP figure survived from FDR correction (q < 0.05). We didn’t perform FDR correction for NSPN and MICs datasets because too few parcels survived. In comparison, for the *t* and asymmetry maps between the different datasets we used non-thresholded maps.

We performed the correlations between brain maps using variogram permutations to test the spatial autocorrelation. The variogram quantifies, as a function of distance *d*, the variance between all pairs of points spatially separated by *d*. Pure white noise, for example, which has equal variation across all spatial scales, has a flat variogram (i.e., no distance dependence). Brain maps with very little spatial autocorrelation will therefore have a variogram that is nearly flat. Strongly autocorrelated brain maps exhibit less variation among spatially proximal regions (at small *d*) than among widely separated regions, and are therefore characterized by positive slopes in their variograms (Burt et al., 2020; Vos de Wael et al., 2020). We obtained the geodesic distance matrix of the left hemisphere from multimodal parcellation (i.e., 180*180) and produced 1000 permuted spatial autocorrelation-preserving surrogate brain maps whose variograms were approximately matched to a target brain map’s variogram.

### Heritability analysis

Referring to our previous work (Bayrak et al., 2022; Valk et al., 2021, 2022; Wan et al., 2022), we analyzed heritability based on the twin design of HCP. Briefly, we calculated the heritability estimates with standard errors via the Sequential Oligogenic Linkage Analysis Routines (SOLAR, version 9.0.0). SOLAR uses maximum likelihood variance decomposition methods to determine the relative importance of familial and environmental influences on a phenotype by modeling the covariance among family members as a function of genetic proximity (Almasy & Blangero, 1998). Heritability, i.e., narrow-sense heritability *h*^2^, represents the proportion of the phenotypic variance (σ^2^ ) accounted for by the total additive genetic variance (σ^2^ ), that is *h*^2^ = σ^2^ /σ^2^ . Phenotypes exhibiting stronger covariances between genetically more similar individuals, than between genetically less similar individuals, have higher heritability. In this study, we quantified the heritability of asymmetry of functional gradients. We added age, sex, age^2^, and age*sex as the covariates to our model.

### Microstructure-function asymmetric coupling

We conducted three approaches to understand the relationship between microstructural and functional asymmetry. We first split the mean microstructural asymmetry map into 10 bins and plotted the FC asymmetry along the 10 bins to test whether group level FC asymmetry was stronger or weaker in asymmetric or non-asymmetric bins. Second, we correlated the microstructural spatial asymmetry of the FC asymmetry per column at the group and individual levels. Pearson correlation coefficients were used to indicate the degree of region-wise coupling. Last, we correlated the population-wise microstructure per region of FC asymmetry between that region and other regions. The coupling matrix was calculated using Pearson correlation coefficients. The columns are microstructural profiles and rows are functional profiles. An affinity matrix was calculated via normalized angle cosine similarity with threshold of top 10% coupling for microstructural and functional profiles separately. Principal component analysis (PCA) was used to decompose the matrix and we extracted the first 10 eigenvectors. Regions having similar coupling profiles were then embedded together along the PC eigenvectors. These steps were done in Python with the package BrainSpace (Vos de Wael et al., 2020).

### Brain-behavioral association

We used canonical correlation analysis to address the multivariate association between microstructural asymmetry and behavioral scores. CCA is a statistical method that finds linear combinations of two random variables so that the correlation between the combined variables is maximized. In practice, CCA has been mainly implemented as a substitute for univariate general linear model to link different modalities, and therefore, is a major and powerful tool in multimodal data fusion. However, the complicated multivariate formulations and obscure capabilities remain obstacles for CCA and its variants to being widely applied. We separately tested brain-behavioral association for language and mental health. Language scores were acquired by reading and picture vocabulary tests in the NIH toolbox. Mental health scores included depression, anxiety, somatic, avoidant, ADHD, and antisocial problems, which were assessed by Adult Self-Report and DSM-Oriented Scale.

The Picture Vocabulary Test is a CAT format measure of general vocabulary knowledge for ages 3-85 and is considered to be a strong measure of crystallized abilities (those abilities that are more dependent upon past learning experiences and are consistent across the lifespan). The participant is presented with an audio recording of a word and four photographic images on the computer screen and is asked to select the picture that most closely matches the meaning of the word. Higher scores indicate higher vocabulary ability. The Reading Test is a CAT format measure of reading decoding skill and of crystallized abilities, those abilities that are generally more dependent upon past learning experiences and consistent across the life span for ages 7-85. The participant is asked to read and pronounce letters and words as accurately as possible. Higher scores indicate better reading ability. Age-adjusted Scale Score: Participant score is normed using the age appropriate band of Toolbox Norming Sample (bands of ages 18-29, or 30-35), where a score of 100 indicates performance that was at the national average and a score of 115 or 85, indicates performance 1 SD above or below the national average for participants’ age band. We used age adjusted language scores in the current study.

The Adult Self-Report is a 126-item self-report questionnaire for adults (ages 18–59) assessing aspects of adaptive functioning and problems. The questionnaire provides scores for the following syndrome scales: anxious/depressed, withdrawn, somatic complaints, thought problems, attention problems, aggressive behavior, rule-breaking behavior, and intrusive behavior. The questionnaire provides scores for the following DSM-oriented scales: depressive problems, anxiety problems, somatic problems, avoidant personality problems, attention deficit/ hyperactivity problems (inattention and hyperactivity/impulsivity subscales), and antisocial personality problems. Additionally, the questionnaire asks about use of the following substances: tobacco, alcohol, and drugs. Items are rated on a 3-point scale: 0-Not True, 1-Somewhat or Sometimes True, 2-Very True or Often True. We used DSM-oriented scale and gender and age adjusted T-scores as mental health scores in the current study.

We also used pseudo-randomised resampling (clustering the same number of twins for each sample) to test the robustness of brain-behavior association and the brain loading patterns. We started by withdrawing 10% to 90% data and iterated these steps 100 times. Therefore, for each percentage data withdrawal, there would be 100 CCA models and correlations to the anterior-posterior asymmetry pattern. We showed the charts with mean and standard error bars for each percentage data withdrawal.

## Data and code availability

The data we used in the current study are available through the holder’s websites, which have been mentioned in the data section. All the scripts and visualization are openly available at a GitHub repository (https://github.com/wanb-psych/microstructural_asymmetry). The packages are completely open for use, see documentations: BigBrainWarp (https://bigbrainwarp.readthedocs.io/en/latest/), BrainStat (https://brainstat.readthedocs.io/en/), BrainSpace (https://brainspace.readthedocs.io/en/latest/), SOLAR (http://solar-eclipse-genetics.org/), and Scikit-learn (https://scikit-learn.org/stable/).

## Acknowledgements

We are grateful to those open datasets including BigBrain, HCP, NSPN, and MICs. BCB acknowledges research support from the National Science and Engineering Research Council of Canada (NSERC Discovery-1304413), Canadian Institutes of Health Research (FDN-154298, PJT-174995), SickKids Foundation (NI17-039), BrainCanada, FRQ-S, and the Tier-2 Canada Research Chairs program. SLV and BCB are furthermore funded by the Helmholtz International BigBrain Analytics and Learning Laboratory (HIBALL), supported by the Helmholtz Association’s Initiative and Networking Fund and the Healthy Brains, Healthy Lives initiative at McGill University. MDH is funded by the German Ministry for Education and Research (BMBF) and the Max Planck Society. SLV is supported by the Otto Hahn Award at Max Planck Society, and BW is supported by the International Max Planck Research School on Neuroscience of Communication: Function, Structure, and Plasticity (IMPRS NeuroCom), Graduate Academy Leipzig, and Mitacs Globalink Research Award.

## Author contributions

Bin Wan: Conceptualization, Methodology, Formal analysis, Writing - Original Draft, Writing - Review & Editing, Visualization, Data curation, Project administration, Funding acquisition.

Amin Saberi: Methodology, Data curation, Writing - Review & Editing

Casey Paquola: Data curation, Writing - Review & Editing

H. Lina Schaare: Methodology, Writing - Review & Editing

Meike D. Hettwer: Data curation, Writing - Review & Editing

Jessica Royer: Data curation, Writing - Review & Editing

Alexandra John: Data curation, Writing - Review & Editing

Lena Dorfschmidt: Data curation, Writing - Review & Editing

Şeyma Bayrak: Data curation, Writing - Review & Editing

Richard A.I. Bethlehem: Writing - Review & Editing

Simon B. Eickhoff: Writing - Review & Editing, Supervision

Boris C. Bernhardt: Data curation, Writing - Review & Editing, Supervision, Funding acquisition

Sofie L. Valk: Conceptualization, Methodology, Data curation, Writing - Original Draft, Writing - Review & Editing, Supervision, Funding acquisition

---

**Supplementary Figure S1.**
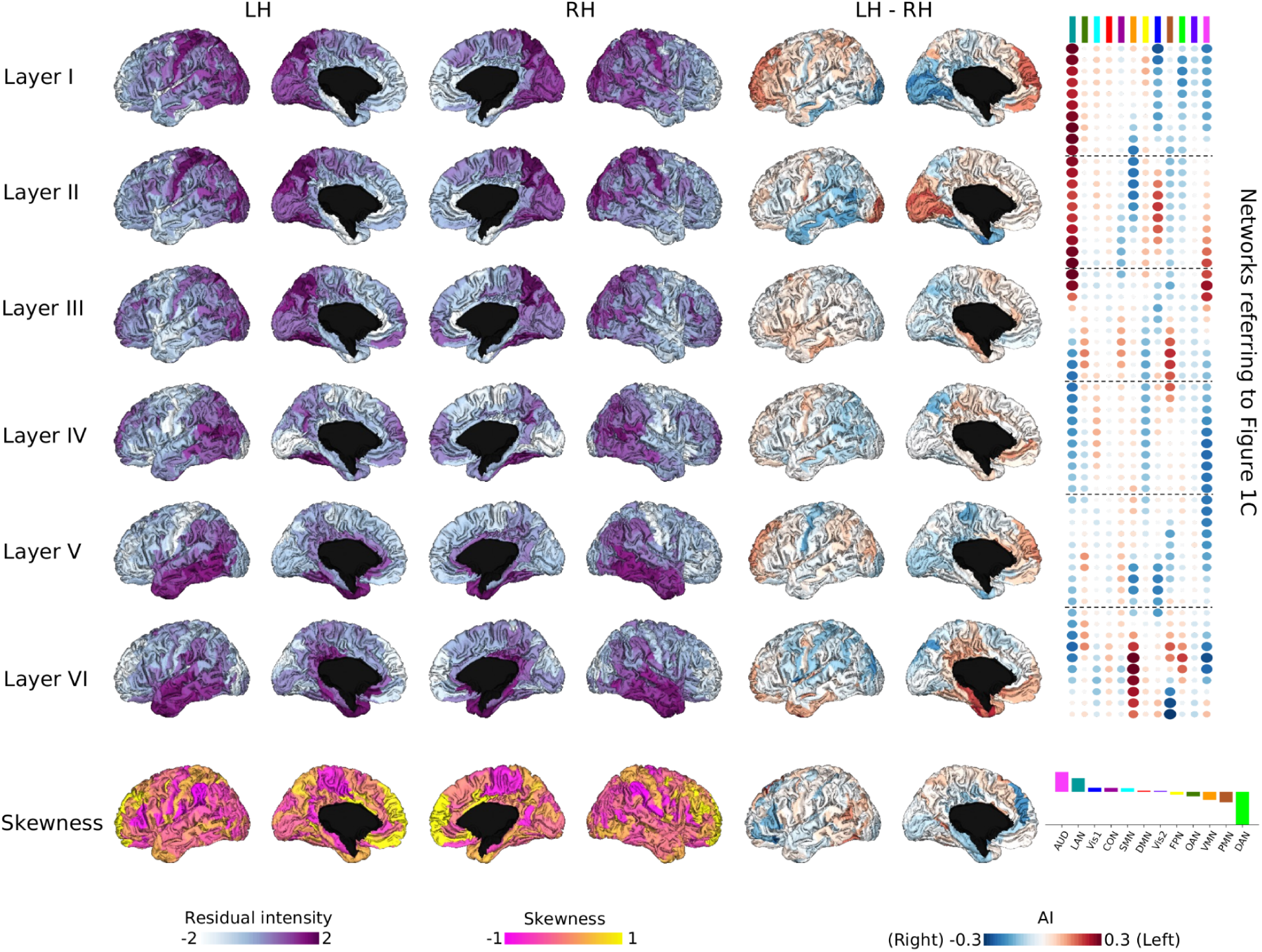
Residual intensity values for each layer.

**Supplementary Figure S2.**
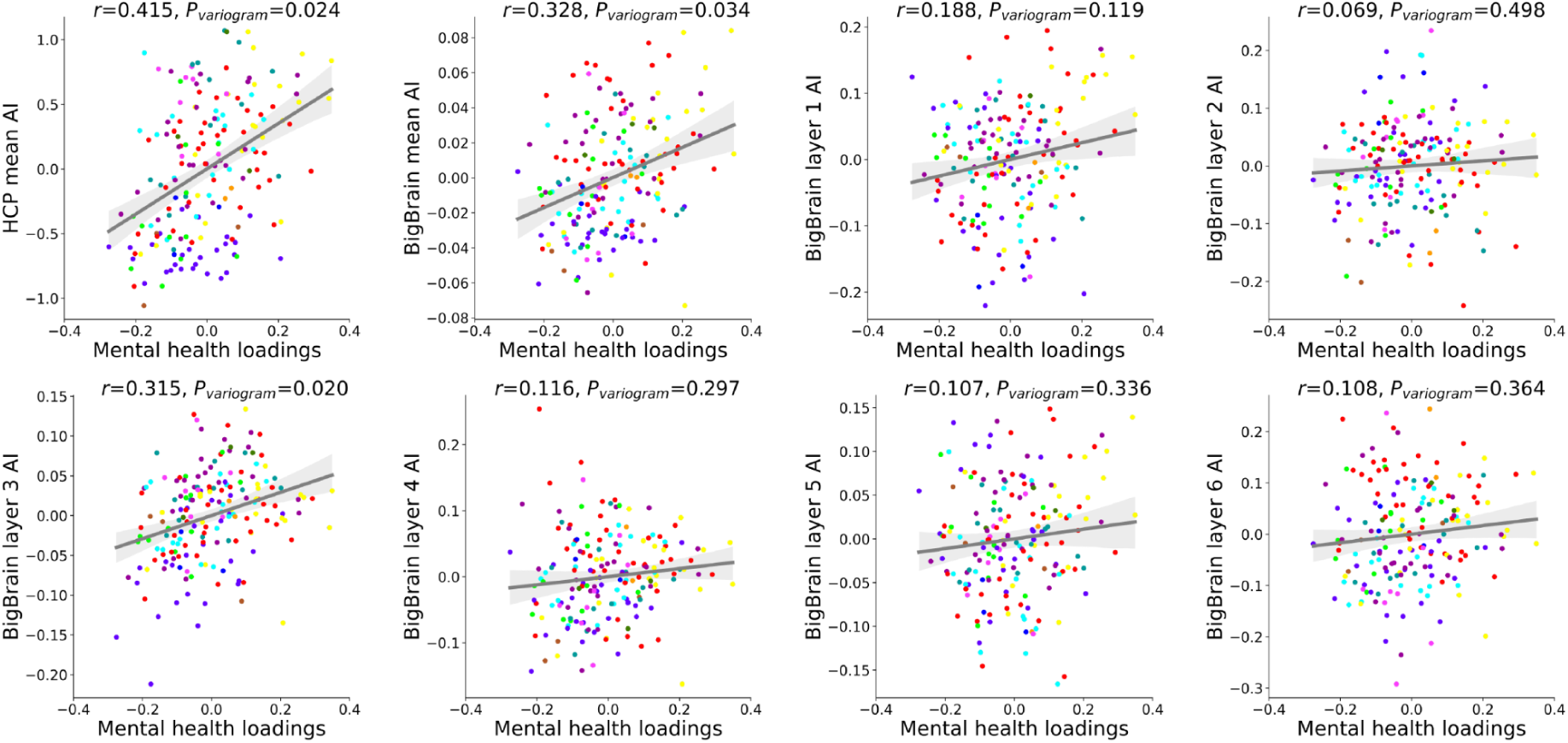
Spatial correlation between CCA mental health brain loadings and AI maps.

**Supplementary Figure S3.**
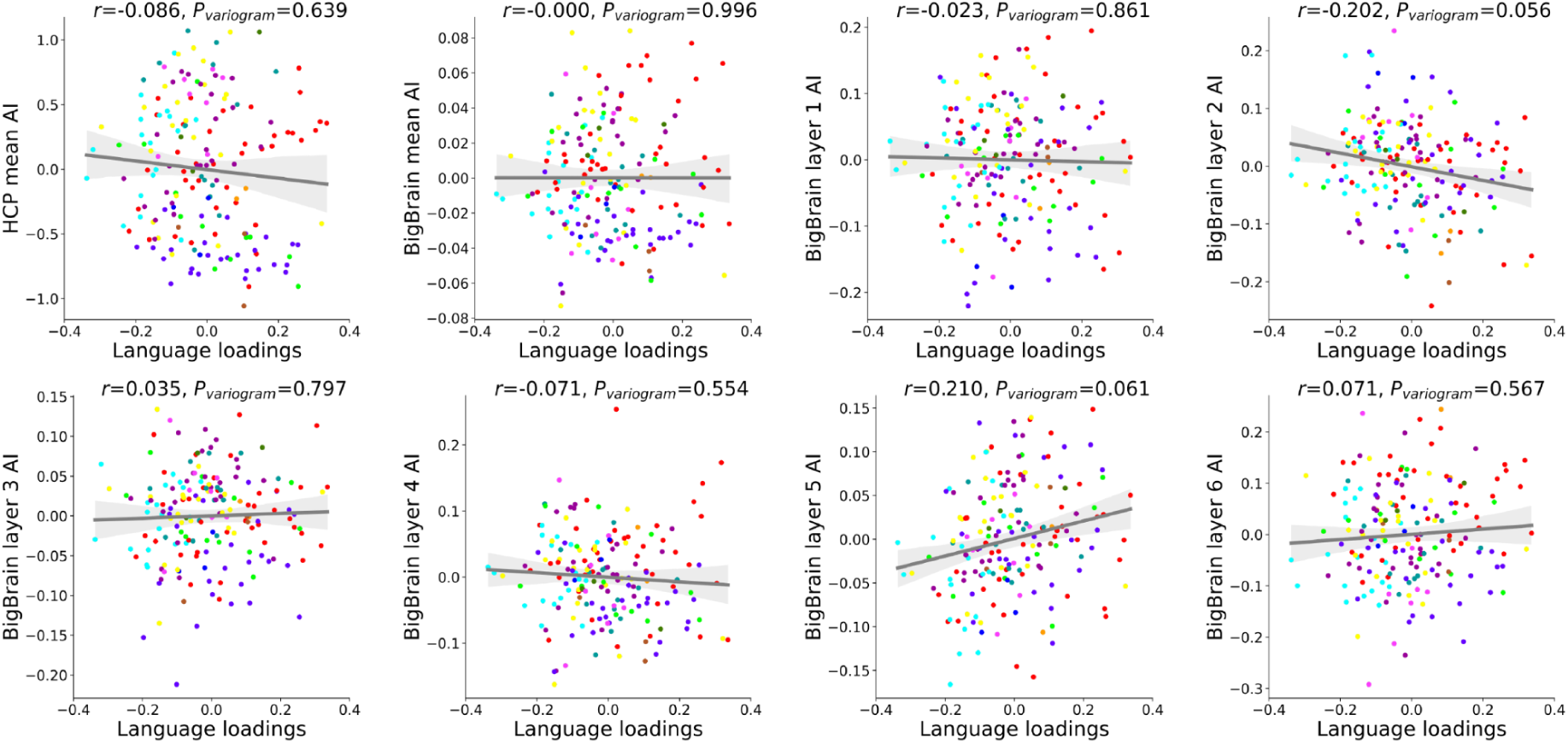
Spatial correlation between CCA language brain loadings and AI maps.

**Supplementary Figure S4.**
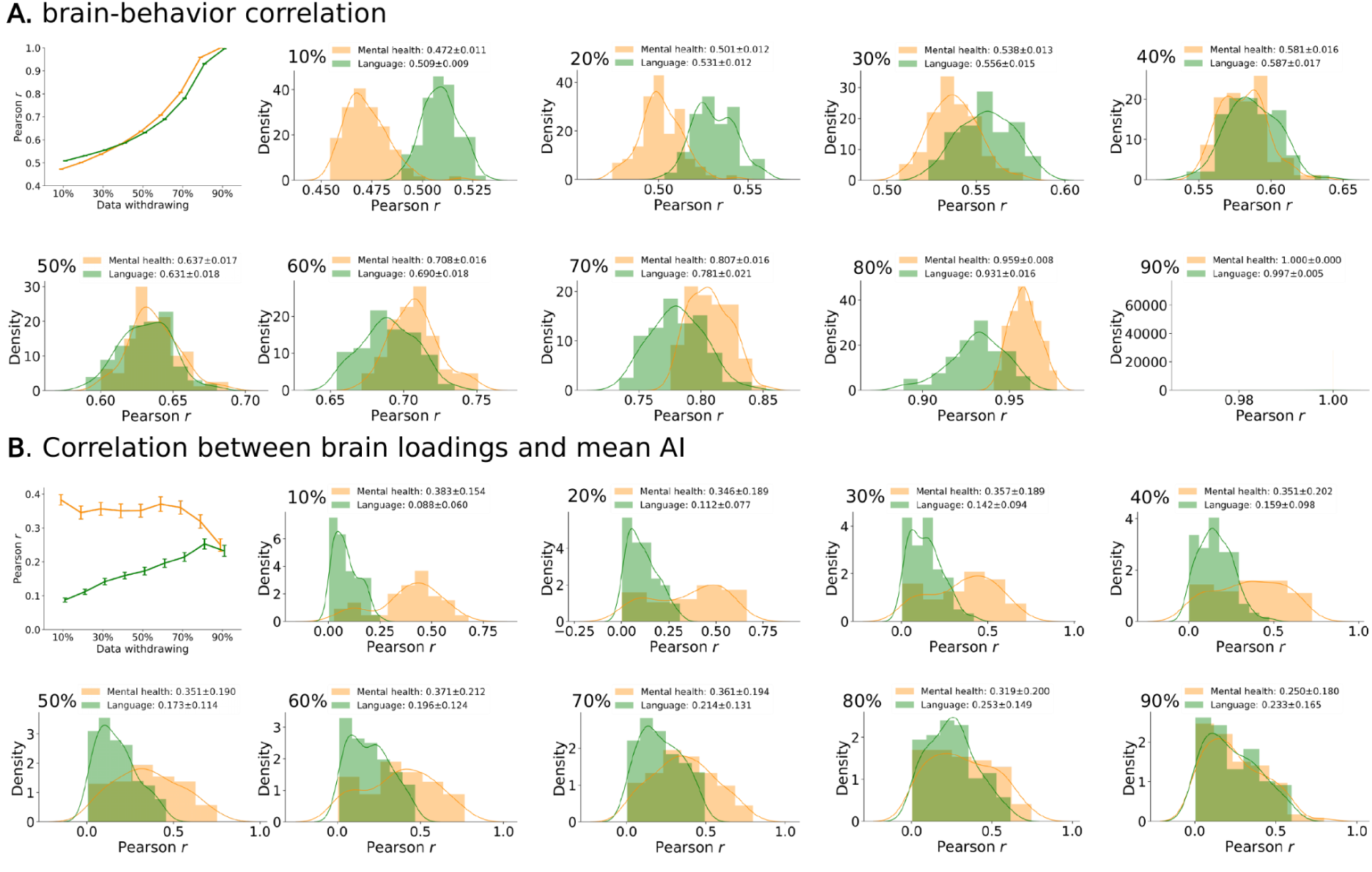
Robustness test for CCA using resampling for 1000 times.

**Supplementary Figure S5.**
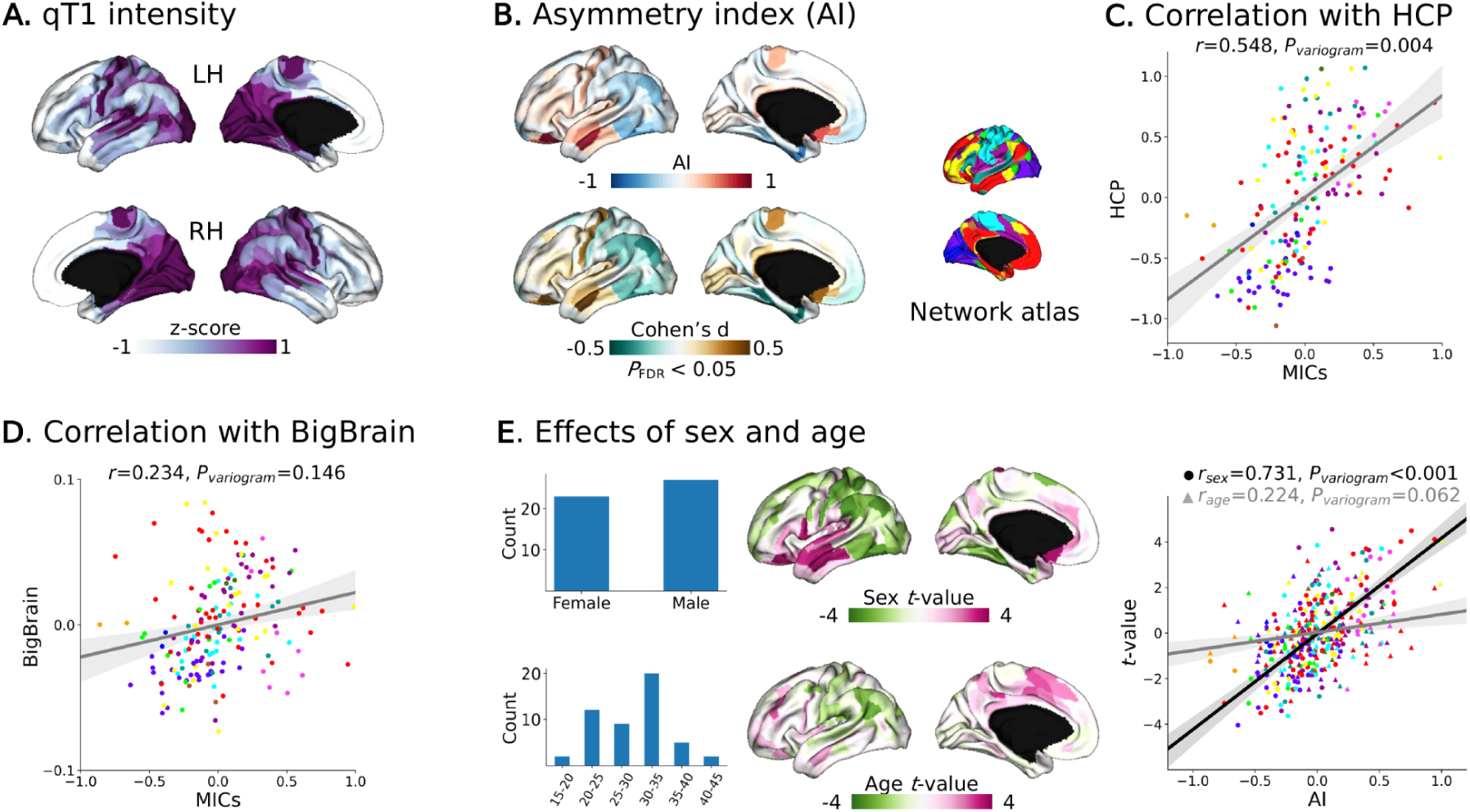
Microstructural asymmetry in MICs.

**Supplementary Figure S6.**
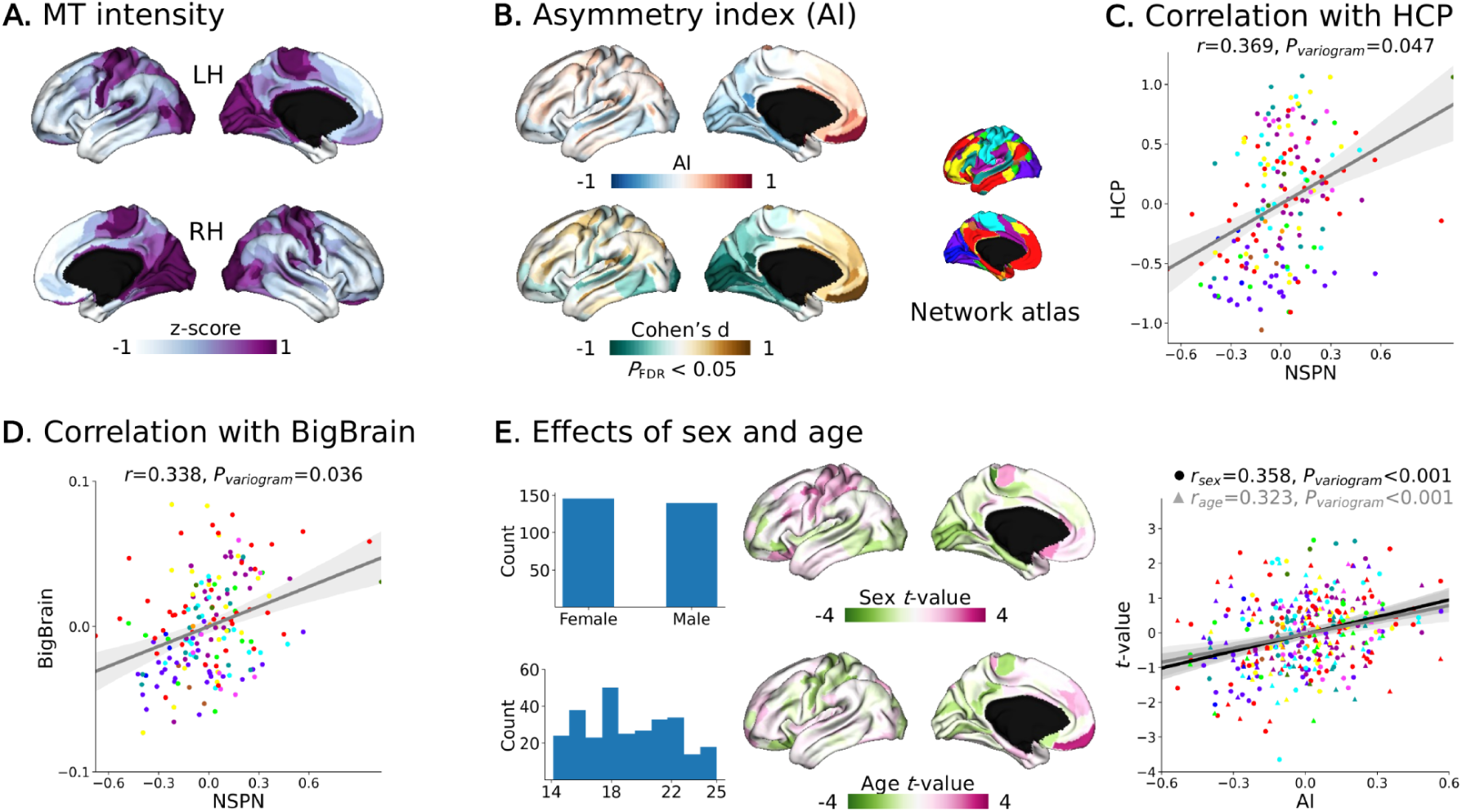
Microstructural asymmetry in NSPN.

## Notes

### Competing Interest Statement

The authors have declared no competing interest.

